# *In utero* and neonatal lipid nanoparticle delivery achieves robust editing in hematopoietic stem cells

**DOI:** 10.1101/2025.05.12.652147

**Authors:** Atesh K. Worthington, Beltran Borges, Elisa Schrader Echeverri, Fareha Moulana Zada, Tony Lum, Andrew Davis, Kun Jia, Barbara Perez, Amanda Graveson, Daniel Oder, Marco A. Cordero, Hyejin Kim, Ryan Zenhausern, Ozgenur Celik, Travis McCreary, Harshmeet Singh, Ramsey Majzoub, Cindy Shaw, Paula Gutierrez-Martinez, Marzhana Omarova, Chris Blanchard, Sean Burns, M. Kyle Cromer, James E. Dahlman, Tippi C. MacKenzie

## Abstract

Efficient delivery of genome editing reagents to hematopoietic stem cells (HSCs) has limited the development of *in vivo* gene editing therapies for hematologic disease. Here, we exploit developmental hematopoiesis to enable HSC targeting using clinically scalable lipid nanoparticles (LNPs). During fetal development, HSCs reside in the liver, a tissue that is efficiently accessed by LNPs. We show that in utero delivery of LNPs carrying Cre recombinase or CRISPR-Cas9 components results in transfection and genome editing of bona fide long-term repopulating HSCs. Edited HSCs maintain multilineage reconstitution capacity following transplantation, demonstrating preserved stem cell function. Comparative studies reveal that both fetal and early neonatal delivery permit HSC editing, with greater efficiency during fetal liver hematopoiesis. We further identify an LNP formulation that enhance HSC targeting and enable robust neonatal HSC editing without antibody-mediated targeting. Finally, combined delivery of Cas9 via LNPs and a repair template via adeno-associated virus in neonatal mice enables *in vivo* homology-directed repair in multiple tissues. Together, these findings establish the perinatal period as a therapeutic window for *in vivo* HSC genome editing and provide a scalable strategy for treating severe early-onset hematologic diseases.

## Introduction

Monogenic hematologic diseases represent a compelling opportunity for curative genome editing therapies. Disorders such as hemoglobinopathies, primary immunodeficiencies and inherited bone marrow failure syndromes could potentially be treated by correcting pathogenic variants in hematopoietic stem cells (HSCs). In several conditions, editing only a fraction of HSCs may be sufficient to achieve therapeutic benefit: partial correction of HSCs in hemoglobinopathies can generate enough healthy erythroid cells to ameliorate disease symptoms^1–3^. Recent clinical advances have demonstrated the feasibility of ex vivo HSC gene editing using viral gene addition and CRISPR-based approaches^4–7^. However, these therapies require harvesting bone marrow stem cells, performing ex vivo manipulation, and transplanting edited cells following conditioning regimens that carry substantial risks and morbidity^8^. In addition, individualized cell manufacturing increases cost and limits accessibility^9,10^. Strategies that enable direct *in vivo* editing of HSCs could overcome these barriers by eliminating the need for cell collection, ex vivo culture and transplant conditioning.

Despite this promise, efficient delivery of genome editing reagents to HSCs *in vivo* remains a major challenge. Lipid nanoparticles (LNPs) have emerged as powerful delivery vehicles for genome editing cargos, enabling transient expression of mRNA and guide RNA without genomic integration and supporting scalable manufacturing^11^, as demonstrated by mRNA vaccines^12,13^ and ongoing genome editing clinical trials^14,15^. However, conventional LNP formulations primarily target hepatocytes^16^ and achieve limited delivery to hematopoietic stem cells in the bone marrow. Recent approaches have attempted to overcome this limitation by decorating LNPs with targeting ligands or antibodies directed toward HSC surface markers^17,18^. While effective in some experimental settings, these strategies may complicate manufacturing and clinical translation.

Developmental hematopoiesis offers an alternative strategy for accessing HSCs *in vivo*. During fetal development, HSCs reside primarily in the fetal liver before migrating to the bone marrow later in gestation^19^. There are also circulating HSCs in both mice and humans during the perinatal period. These developmental dynamics may create a unique window during which HSCs are more accessible to systemically delivered therapeutics. Importantly, fetal and neonatal HSCs are highly proliferative^20,21^, a feature that may favor CRISPR/Cas9–mediated genome editing^22^.

Recent advances in prenatal diagnosis and fetal therapy further support the rationale and clinical potential of genome editing during early development. Many severe genetic disorders are now diagnosed before birth through carrier screening, chorionic villus sampling, amniocentesis, and noninvasive prenatal testing^23^. In parallel, clinical experience with fetal interventions such as intrauterine transfusions^24^, stem cell transplantation^25–27^ and enzyme replacement therapies^28,29^ demonstrate the feasibility of delivering therapeutics directly to the fetus. Early postnatal genomic correction has been demonstrated in a child with a urea cycle disorder, suggesting a strategy of prenatal diagnosis and neonatal correction in other settings^15^. Early correction of disease-causing mutations may be particularly valuable for severe congenital hematologic disorders, in which pathology begins before or shortly after birth and correction in childhood involves the morbidity of a bone marrow transplant^30^. The main limitation for clinical applications of this strategy is delivery of LNPs to HSCs.

We hypothesized that LNPs, which naturally accumulate in the liver after systemic delivery, could be used to access hematopoietic stem cells during fetal liver hematopoiesis or in the systemic circulation immediately after birth^31^. In this study, we investigate the biodistribution, HSC transfection efficiency, and genome editing outcomes of LNP delivery during fetal and neonatal stages in mice. We demonstrate that liver-tropic LNPs can access and edit bona fide long-term repopulating HSCs *in vivo*, identify an LNP formulation that improves neonatal HSC targeting, and establish the feasibility of *in vivo* homology-directed repair using combined LNP and adeno-associated virus (AAV) delivery. These results identify the perinatal period as a therapeutic window for *in vivo* HSC genome editing and support the development of scalable genome editing strategies for early-onset hematologic diseases.

## Results

### *In utero* delivery of LNPs results in broad fetal tissue transfection and fetal HSC targeting

To evaluate LNP delivery during early development, we first tested LNP^67^, a formulation previously reported to deliver mRNA to adult bone marrow (BM) HSCs^32^. We injected fetal mTmG^33^ reporter mice, in which Cre recombinase induces a fluorescent switch from tdTomato to GFP, via intrahepatic injection at embryonic day (E) 13.5–14.5 with 1.5–3 mg/kg LNP^67^ encapsulating Cre mRNA (**Fig. 1A**). Tissues were harvested 6–8 days later to assess biodistribution.

**Figure 1.**
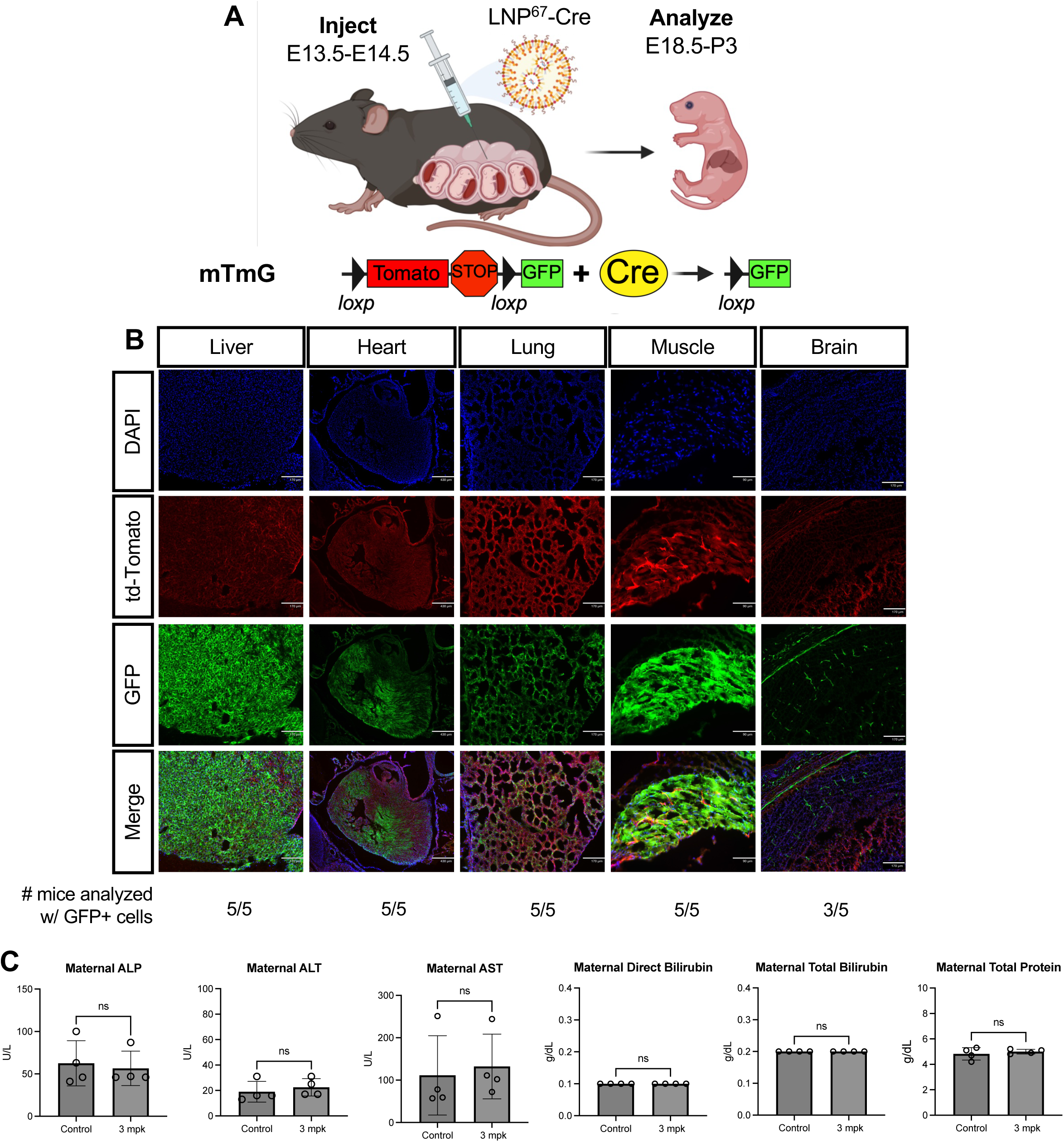
Wide biodistribution following *in utero* delivery of LNP packaged with Cre- recombinase mRNA. **A)** Experiment schematic. E13.5-E14.5 mTmG mice were injected intrahepatically with LNP^67^- Cre and tissues were harvested 6-8 days later. **B)** Representative images of GFP (green) and tdTomato expression (red) in liver, heart, lung, muscle and brain. **C)** Maternal serum liver function tests 5-6 days post PBS or 3 mpk LNP^67^-Cre *in utero* fetal injection. Data are mean+/-SEM; each dot represents one dam, PBS n=5, 3 mpk LNP^67^-Cre n=4. Differences were analyzed Mann-Whitney test, P=NS

Fluorescence microscopy revealed robust GFP expression in the liver, heart, lung and skeletal muscle of all injected mice, with detectable GFP expression in the brain in a subset of animals (3 of 5) (**Fig. 1B**), demonstrating widespread fetal tissue transfection. Because fetal therapy involves exposure of both the fetus and the pregnant dam, we next assessed maternal safety by measuring maternal serum liver function tests 6 days after injection. We observed no significant differences in alanine aminotransferase (ALT), aspartate aminotransferase (AST), alkaline phosphatase (ALP), bilirubin or total protein between PBS- and LNP-injected dams (**Fig. 1C**).

We next asked whether LNP^67^ could transfect hematopoietic stem cells during fetal development. Because the fetal liver serves as the primary site of hematopoiesis at this stage, we quantified GFP⁺ HSCs in fetal livers by flow cytometry (**Fig. 2A**). We detected GFP expression in a subset of immunophenotypic HSCs, although at lower frequencies than in non-hematopoietic liver cells (**Fig. 2B-D**).

**Figure 2.**
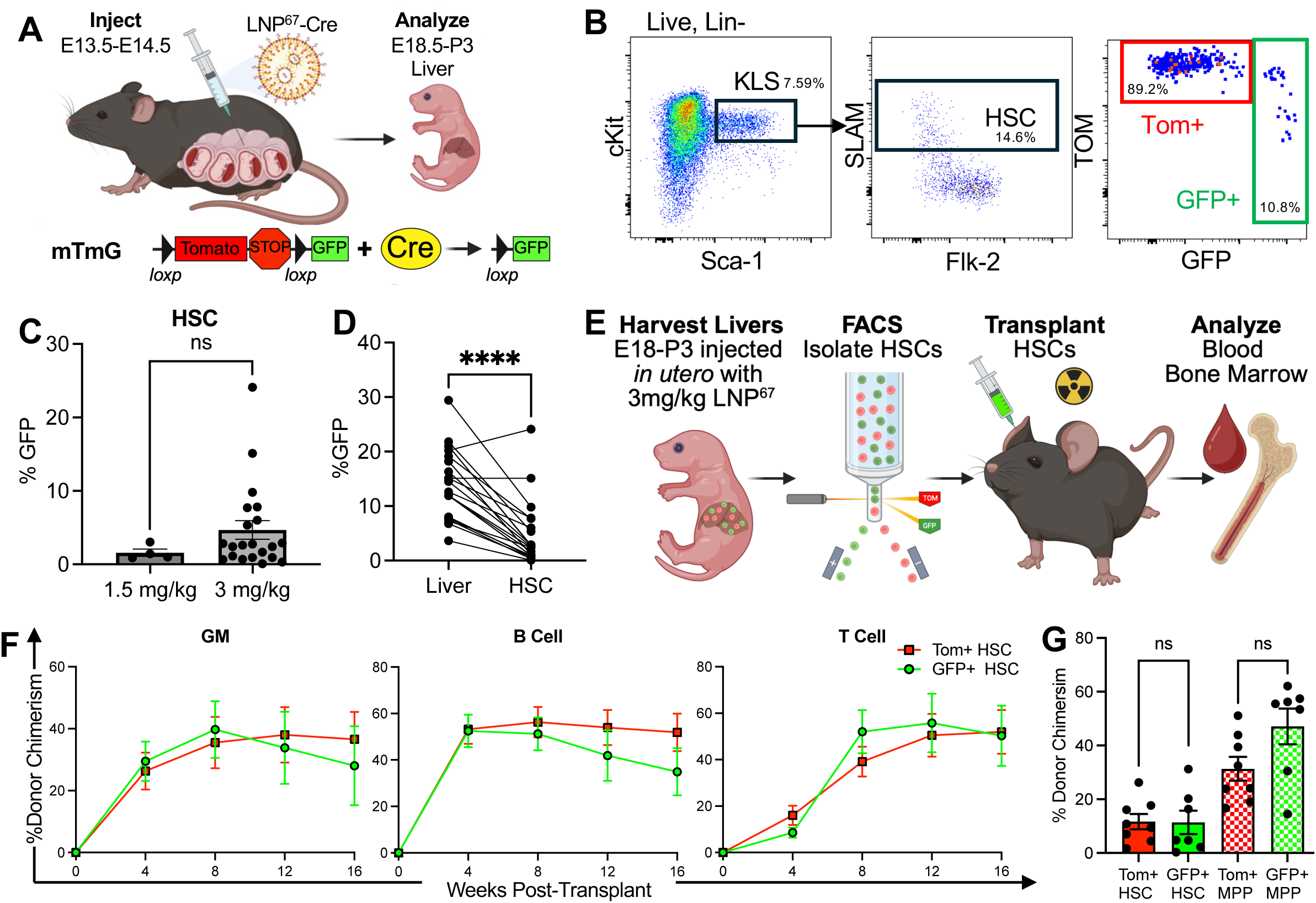
*In utero* LNP-Cre delivery results in transfection of *bona fide* HSC. **A)** Experiment schematic. **B)** Gating strategy to identify LNP-transfected HSCs (Lin- cKit+ Sca+ SLAM+ GFP+). **C)** % GFP+ liver HSCs. 1.5 mg/kg (n=4), 3 mg/kg (n=21). P= not significant (NS) (Student’s t-test). **D)** Percentage of GFP+ liver cells vs. HSCs in livers of LNP-injected mice (3 mg/kg, n=21). ****P<0.0001 (Paired student’s t-test). **E)** Experiment schematic of HSC transplants after isolating GFP+ HSCs as presented in 1C. **F)** % GFP+ (green circle, n=7) and Tom+ (red squares, n=8) granulocyte and macrophages (GM), B cells and T cells in blood of transplanted mice weeks post-transplant. P=NS between mean GFP+ vs Tom+ at each time point by Student’s t-test. Data are mean+/-SEM; at least 2 independent experiments. **G)** Percentage of donor-derived HSCs (solid bars) and multipotent progenitors (MPPs, patterned bars, Lin-cKit+Sca+SLAM-Flk2+) in BM of GFP+ (green bars) and Tom+ (red bars) HSC transplanted mice 16+ weeks post-transplant. P=NS between mean GFP+ vs Tom+ HSC or MPP (Student’s t-test). Data are mean+/-SEM; each dot represents an individual animal, least 2 independent experiments.

To determine whether these transfected immunophenotypic HSCs represented functional long-term repopulating stem cells, we performed transplantation assays. GFP⁺ liver HSCs were isolated 4–5 days after LNP^67^ injection and transplanted into sublethally irradiated wild-type recipients alongside control tdTomato⁺ HSCs from uninjected littermates (**Fig. 2E**). Donor-derived cells were readily detected in the peripheral blood of recipient mice and contributed to granulocyte-macrophage, B cell and T cell lineages at levels comparable to control HSCs (**Fig. 2F and Fig. S1A**). In addition, donor-derived HSCs and multipotent progenitors were detected in the bone marrow of recipients more than 16 weeks after transplantation (**Fig. 2G** and **Fig. S1B**). These findings demonstrate that LNP^67^-mediated delivery can achieve transfection in bona fide long-term repopulating HSCs without impairing their capacity for multilineage hematopoietic reconstitution.

### Developmental timing influences tissue distribution of LNP-mediated genome editing

Having established that LNPs can transfect fetal HSCs, we next asked whether LNPs could deliver CRISPR–Cas9 genome editing cargos during fetal and neonatal stages. To address this question, we used Traffic Light Reporter (TLR^34^) mice, in which repair of Cas9-induced double-stranded breaks through insertions or deletions can restore the reading frame of an RFP reporter (**Fig. 3A**). This system enables quantification of genome editing by detection of RFP- positive cells.

**Figure 3.**
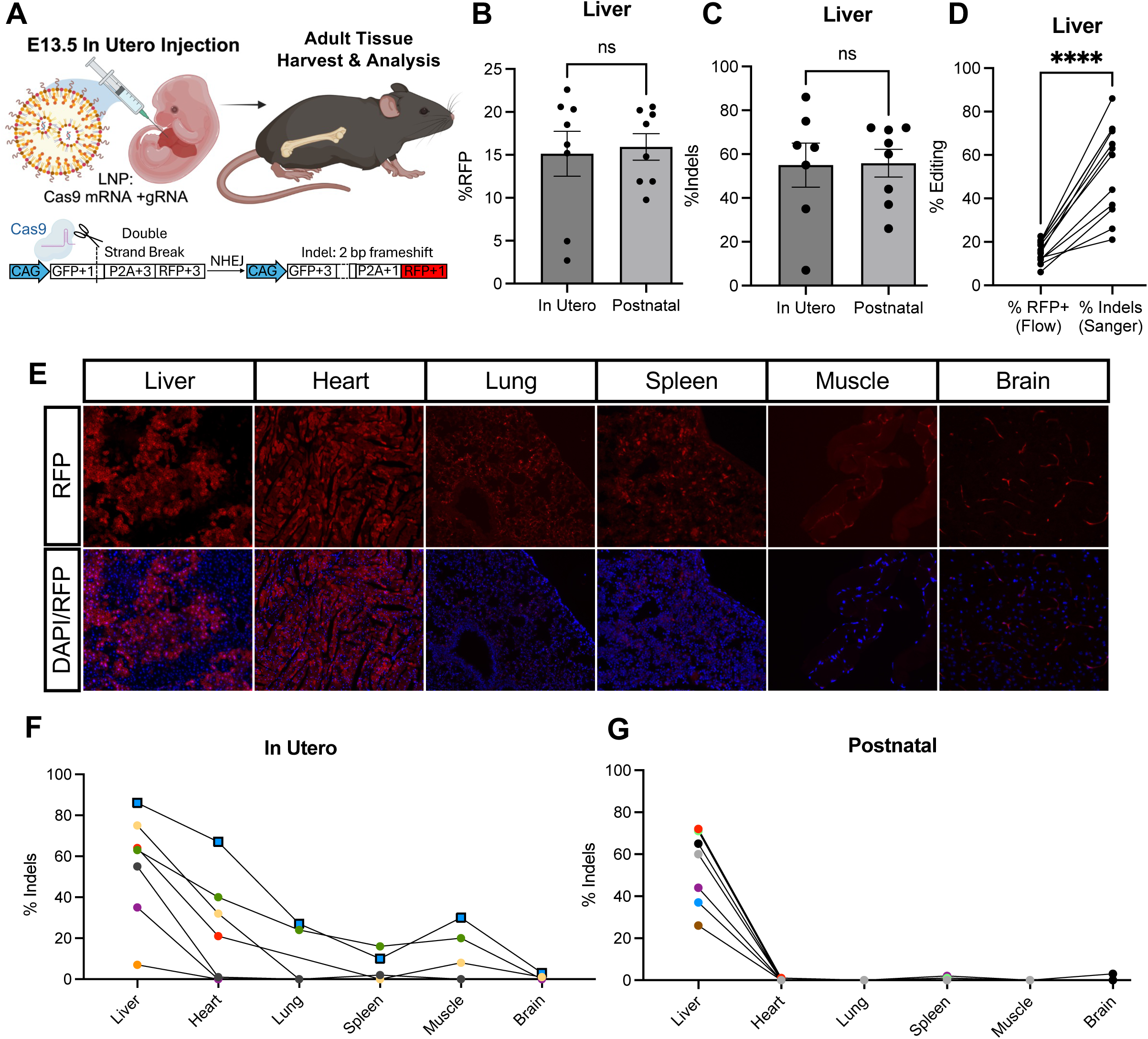
Comparative analysis of genome editing following *in utero* vs neonatal LNP delivery. **A)** Experiment schematic. E13.5 and P0 TLR mice were injected intrahepatically with LNP- Cas9/gRNA. Tissues were harvested in adult mice. **B)** Percentage of RFP+ cells by flow cytometry in the liver mice injected *in utero* (n=8) or postnatally (n=8) with 3 mg/kg of LNP. P=NS (Mann-Whitney test). **C)** Percentage of InDels by Sanger sequencing of cells in the liver mice injected *in utero* (n=8) or postnatally (n=8) with 3 mg/kg of LNP. P=NS (Mann-Whitney test). **D)** Comparison of % editing in liver cells quantified by RFP expression (by flow) or InDel frequency (by Sanger sequencing). ****P<0.0001 (Paired student’s t-test). **E)** Representative images of RFP (red) expression in liver, heart, lung, spleen, muscle and brain. **F)** Percentage of InDels by Sanger sequencing of cells in the liver, heart, lung, spleen, muscle and >brain mice injected *in utero* (n=8) 3 mg/kg of LNP. Each dot represents an individual animal. Lines connect data from the same mouse. Square symbols represent the mouse that the representative images in E come from. or postnatally (n=8) with 3 mg/kg of LNP. P=NS (Mann-Whitney test). **G)** Percentage of InDels by Sanger sequencing of cells in the liver, heart, lung, spleen, muscle and brain mice injected postnatally (n=8) 3 mg/kg of LNP. Each dot represents an individual animal. Lines connect data from the same mouse. Data are mean+/-SEM; each dot represents an individual animal, least 3 independent experiments.

We injected fetal TLR mice intrahepatically at embryonic day (E) 13.5 and neonatal (P0) mice via the temporal vein with 3 mg/kg of a standard liver-tropic LNP formulation encapsulating Cas9 mRNA and a TLR-targeting guide RNA. Mice were analyzed as adults (8-12 weeks post injection) to assess durable editing outcomes.

Flow cytometry revealed comparable frequencies of RFP⁺ cells in the livers of mice treated in utero or postnatally (**Fig. 3B**). Quantification of editing by Sanger sequencing similarly demonstrated comparable levels of insertions and deletions (InDels) between the two delivery timepoints (**Fig. 3C**). Notably, indel frequencies measured by sequencing were substantially higher than RFP expression detected by flow cytometry, consistent with the fact that only a subset of non-homologous end joining (NHEJ) events restore the correct reading frame required for RFP expression (**Fig. 3D**).

We next examined editing across additional tissues. Following in utero delivery, we detected robust editing in multiple organs (including heart, lung, spleen and skeletal muscle) in addition to the liver (**Fig. 3E**). In contrast, neonatal delivery resulted in editing largely restricted to the liver, with minimal or undetectable editing in extrahepatic tissues (**Fig. 3F**). Confocal imaging confirmed widespread RFP expression in several tissues following fetal delivery, with particularly strong editing in the liver and heart (**Fig. 3G**). Thus, LNP-mediated genome editing is efficient during both fetal and neonatal stages, but that fetal delivery enables substantially broader extrahepatic editing across multiple tissues.

### *In vivo* LNP delivery edits functional long-term HSCs

We next asked whether LNP-mediated genome editing during fetal or neonatal stages results in editing of HSCs that persist into adulthood. To address this, we analyzed bone marrow from adult TLR mice that had received liver-tropic LNP-Cas9/gRNA either in utero (E13.5) or shortly after birth (P0). Edited cells were identified by RFP expression via flow cytometry (**Fig. 4A**). We observed robust HSC editing after *in utero* LNP injection; an average of 14% overall and up to 20-36% in some animals (**Fig. 4B, C**). Interestingly, an average of 4% HSC editing was also detected after neonatal injection, although at significantly lower levels compared to *in utero* injection animals (**Fig. 4C**), suggesting that the time window for accessing HSCs extends beyond the prenatal period (despite overall inefficient editing in other extrahepatic tissues (**Fig. 3F**)).

**Figure 4.**
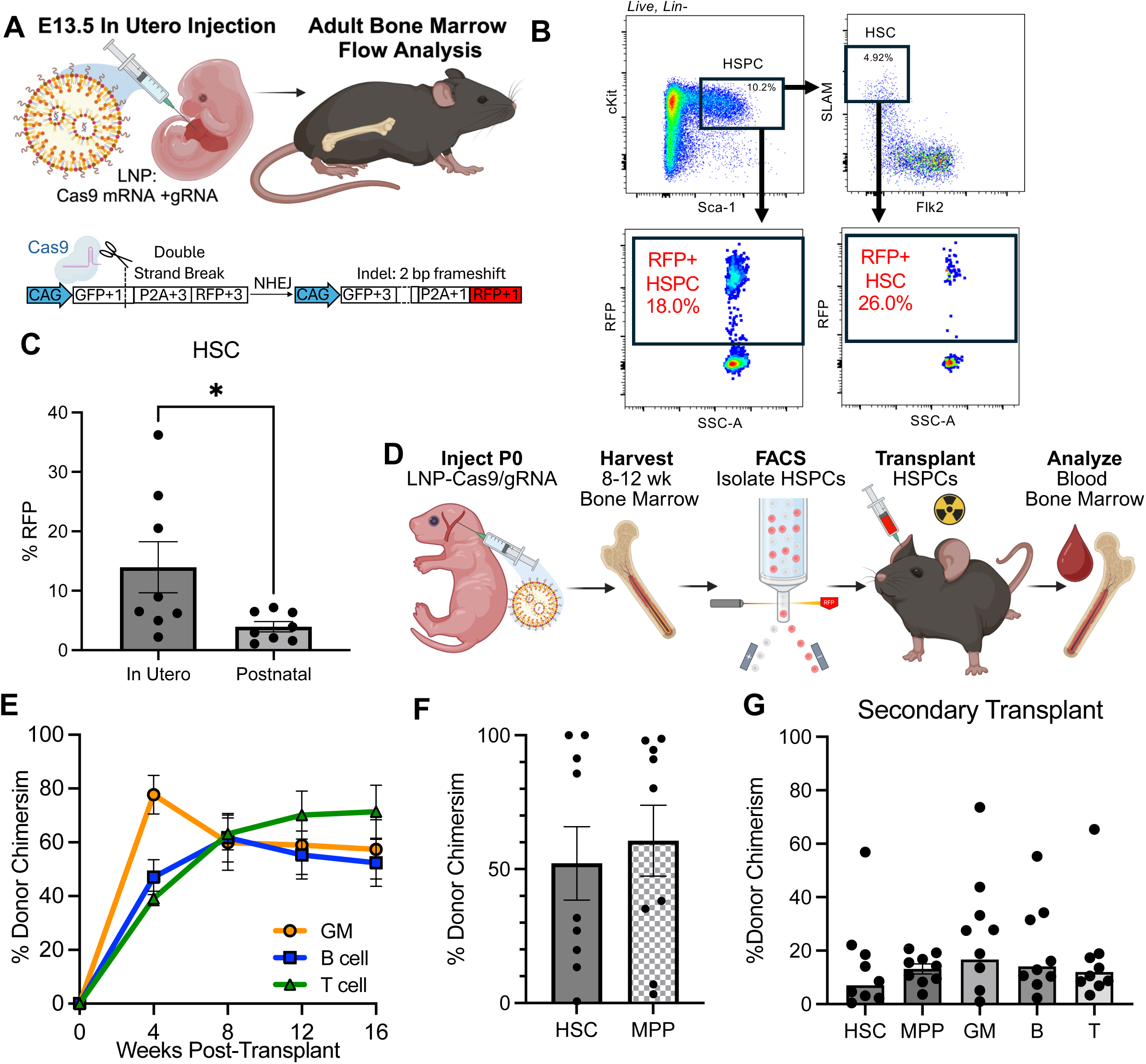
*In utero* or neonatal injection of liver tropic LNPs carrying Cas9/gRNA results in editing of HSCs. **A)** Experiment schematic. E13.5 and P0 TLR mice were injected intrahepatically with LNP- Cas9/gRNA. BM and livers were harvested in adult mice. **B)** Gating strategy to identify Cas9-edited HSPC (KLS: Lin- cKit+ Sca+ RFP+) and HSCs (Lin- cKit+ Sca+ SLAMmid/hi Flk2- RFP+). **C)** Percentage of RFP+ HSCs in BM of mice injected *in utero* (n=8) or postnatally (n=8) with 3 mg/kg of LNP. *P<0.05 (Mann-Whitney test). **D)** Experiment schematic of HSPC transplants. RFP+ HSPCs were isolated from postnatally injected animals presented in 4C. **E)** Percentage of RFP+ (n=10) GM, B cells and T cells in blood of transplanted mice weeks post transplant. **F)** Percentage of donor-derived HSCs (solid bars) and MPPs (patterned bars) in BM of RFP+ HSPC transplanted mice 16+ weeks post-transplant. **G)** Percentage of donor-derived HSCs, MPPs in BM and GM, B and T cells in peripheral blood of secondary transplanted mice 16+ weeks post-transplant.

To determine whether edited cells retained functional stem cell properties, we performed transplantation assays. RFP⁺ HSPCs were isolated from the bone marrow of treated mice and transplanted into sublethally irradiated wild-type recipients (**Fig. 4D**). Donor-derived cells were readily detected in peripheral blood and contributed to granulocyte-macrophage, B cell and T cell lineages over time, demonstrating durable multilineage reconstitution (**Fig. 4E** and **Fig. S2A**). In addition, donor-derived HSCs and multipotent progenitors were detected in the bone marrow of recipient mice more than 16 weeks after transplantation (**Fig. 4F** and **Fig. S2B**).

As a more stringent test of stem cell function, we performed secondary transplantation by transferring whole bone marrow from primary recipients into lethally irradiated secondary recipients. Donor-derived granulocyte-macrophage, B cell and T cell populations were detected in peripheral blood, and donor-derived HSCs and multipotent progenitors were detected in the bone marrow of secondary recipients more than six months after transplantation (**Fig. 4G**). These findings confirm that LNP-mediated genome editing occurs in bona fide long-term repopulating hematopoietic stem cells and does not impair their capacity for long-term multilineage hematopoietic reconstitution.

### Engineered LNP formulation enhance neonatal editing of hematopoietic stem cells

Although liver-tropic LNPs enabled editing of HSCs following neonatal delivery, editing efficiency in HSCs remained lower than that observed following fetal delivery. We therefore asked whether modifying the ionizable lipid component of the LNP formulation could improve HSC targeting during neonatal administration. We evaluated an alternative LNP formulation containing distinct ionizable lipids that were predicted to alter biodistribution and cellular uptake. Neonatal TLR mice were injected via the temporal vein at postnatal day 0 (P0) with 3 mg/kg of LNP-Cas9/gRNA using either the standard liver-tropic formulation or the modified, HSC-tropic formulation (**Fig. 5A**). Mice were analyzed as adults to assess long-term editing outcomes.

**Figure 5.**
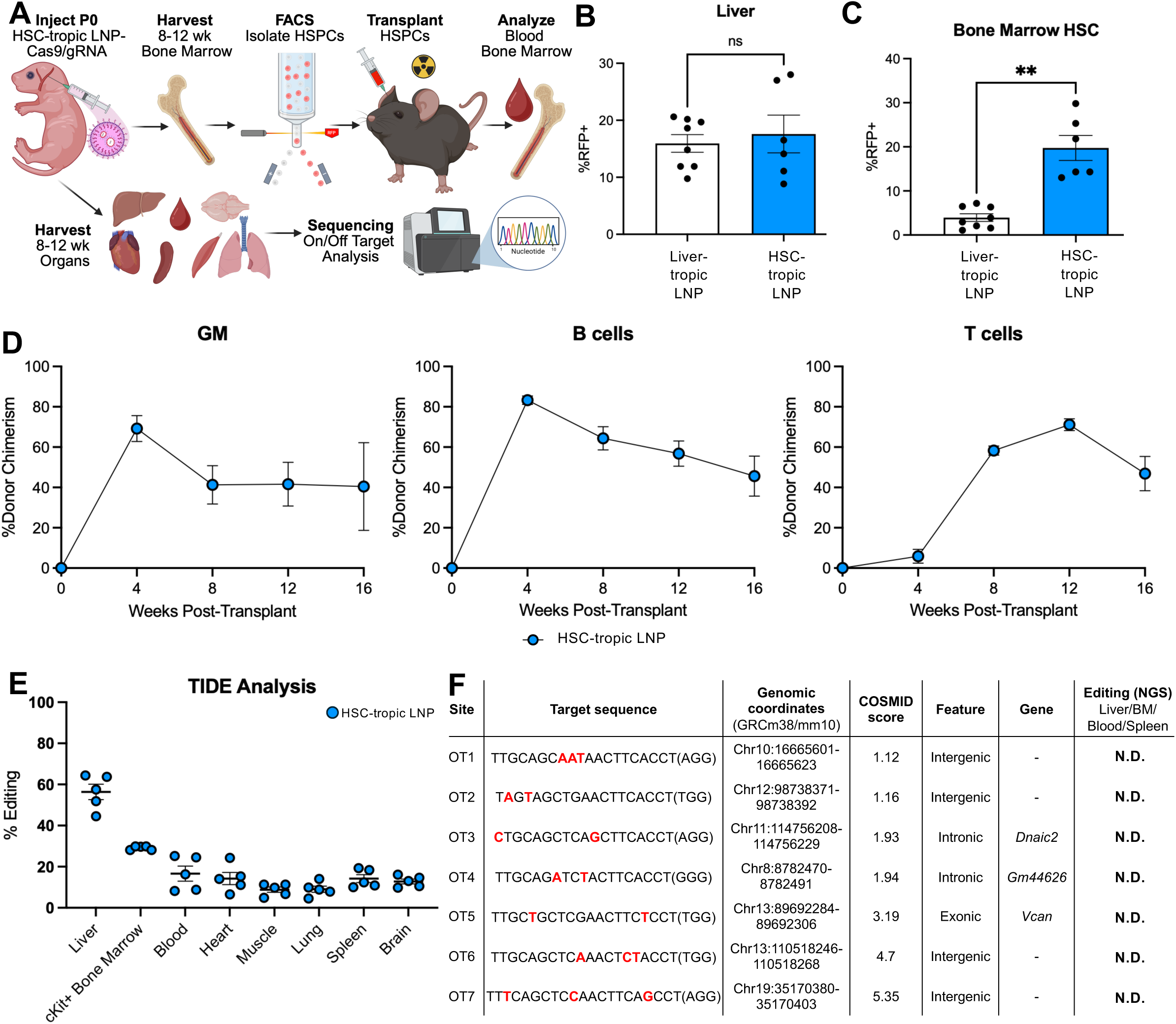
Neonatal injection of HSC-tropic LNPs carrying Cas9/gRNA results in robust editing of HSCs and no observed off-target editing. **A)** Experiment schematic. P0 TLR mice were injected via the temporal vein with HSC-tropic LNP-Cas9/gRNA. Tissues were harvested in adult mice for analysis and HSPCs were isolated for transplant into sublethally irradiated non-fluorescent mice via FACS. **B)** Percentage of RFP+ cells by flow cytometry in the liver of mice injected postnatally with 3 mg/kg of different LNP formulations. P=NS (One-way ANOVA with Tukey’s multiple comparisons test). **C)** Percentage of RFP+ of HSCs in the bone marrow by flow cytometry of mice injected postnatally with 3 mg/kg of different LNP formulations. P=NS (One-way ANOVA with Tukey’s multiple comparisons test). **P<0.01, ***P<0.001 **D)** Percent donor chimerism (HSC-tropic #1: blue circles n =8, HSC-tropic #2: pink square n=7) granulocyte and macrophages (GM), B cells and T cells in blood of transplanted mice weeks post-transplant. P=NS between mean HSC-tropic #1 and #2 at each time point by Student’s t- test. Data are mean+/-SEM; at least 2 independent experiments. **E)** On target editing rates from various tissues of mice treated with HSC-tropic LNP #1 (blue circles, n=5) and HSC-tropic LNP #2 (pink squares, n=3) of 2 LNPs, determined by analysis of Sanger traces on TIDE. Each dot/square is an individual mouse, data are mean+/-SEM **F)** Surveyed off-target sites, selected using COSMID (*in silico*), DISCOVER-seq, and GUIDE- seq (empirical), and observed off-target editing rates in liver, cKit+ bone marrow, blood and spleen as measured by NGS and analyzed by CRISPResso; N.D.= not detected.

Flow cytometry revealed similar levels of editing in liver cells across both formulations, indicating that the modified LNPs retained efficient hepatic delivery (**Fig. 5B** and **Fig. S3A**). In contrast, analysis of bone marrow demonstrated a marked increase in editing of hematopoietic stem cells following delivery of the modified formulation compared with the standard liver-tropic LNP (**Fig. 5C**). These results indicate that altering the ionizable lipid component of the LNP formulation can substantially improve HSC targeting during neonatal delivery.

To determine whether edited cells retained functional stem cell properties, we isolated RFP⁺ HSPCs from the bone marrow of neonatally treated mice and transplanted them into sublethally irradiated wild-type recipients. Donor-derived cells contributed to granulocyte-macrophage, B cell and T cell lineages in peripheral blood over time, demonstrating durable multilineage hematopoietic reconstitution (**Fig. 5D** and **Fig. S3B**). Thus, genome editing using the optimized LNP formulation does not impair HSC function.

We next examined editing in other tissues. Sanger sequencing revealed detectable editing in several tissues (including heart and skeletal muscle) following delivery of the modified HSC- tropic formulation (**Fig. 5E**), whereas editing after neonatal delivery of the standard liver-tropic LNP was largely restricted to the liver (**Fig. 3F**). Thus, modifying LNP composition can extend extrahepatic genome editing beyond the fetal period into other tissues that are impacted by genetic diseases.

Finally, we evaluated the potential for off-target genome editing. Candidate off-target sites were identified using COSMID as well as DISCOVER-seq and GUIDE-seq datasets. Targeted amplicon sequencing of the top seven predicted off-target loci in liver, cKit-enriched bone marrow, blood and spleen were analyzed. We detected no editing above the assay’s limit of detection at any of the seven candidate off-target loci analyzed (**Fig. 5F**). Our findings demonstrate that optimized LNP formulations can substantially enhance neonatal HSC editing while preserving stem cell function and maintaining high editing specificity.

### Combined LNP and AAV delivery enables *in vivo* homology-directed repair

While fortuitous InDels have had great success in the clinic to treat sickle cell anemia and beta-thalassemia, diseases that occur due to deletions or nucleotide changes may require different strategies for correction. Repairing large sections of DNA via homology-directed repair (HDR) is a particularly attractive solution, since the same editing strategy (and therefore drug product) could be used for many patients with different causative variants. However, achieving efficient *in vivo* HDR has been challenging. Having established that optimized an LNP formulation can efficiently deliver Cas9 and guide RNA *in vivo*, we next asked whether this platform could support HDR by co-delivering a DNA repair template.

To quantify HDR *in vivo*, we again used the Traffic Light Reporter (TLR) mouse model. In these mice, a truncated GFP gene can be restored through HDR following a Cas9-induced double-strand break using a donor repair template (**Fig. 6A**). Successful HDR restores GFP expression, whereas NHEJ events generate RFP expression, enabling simultaneous detection of both editing outcomes.

**Figure 6.**
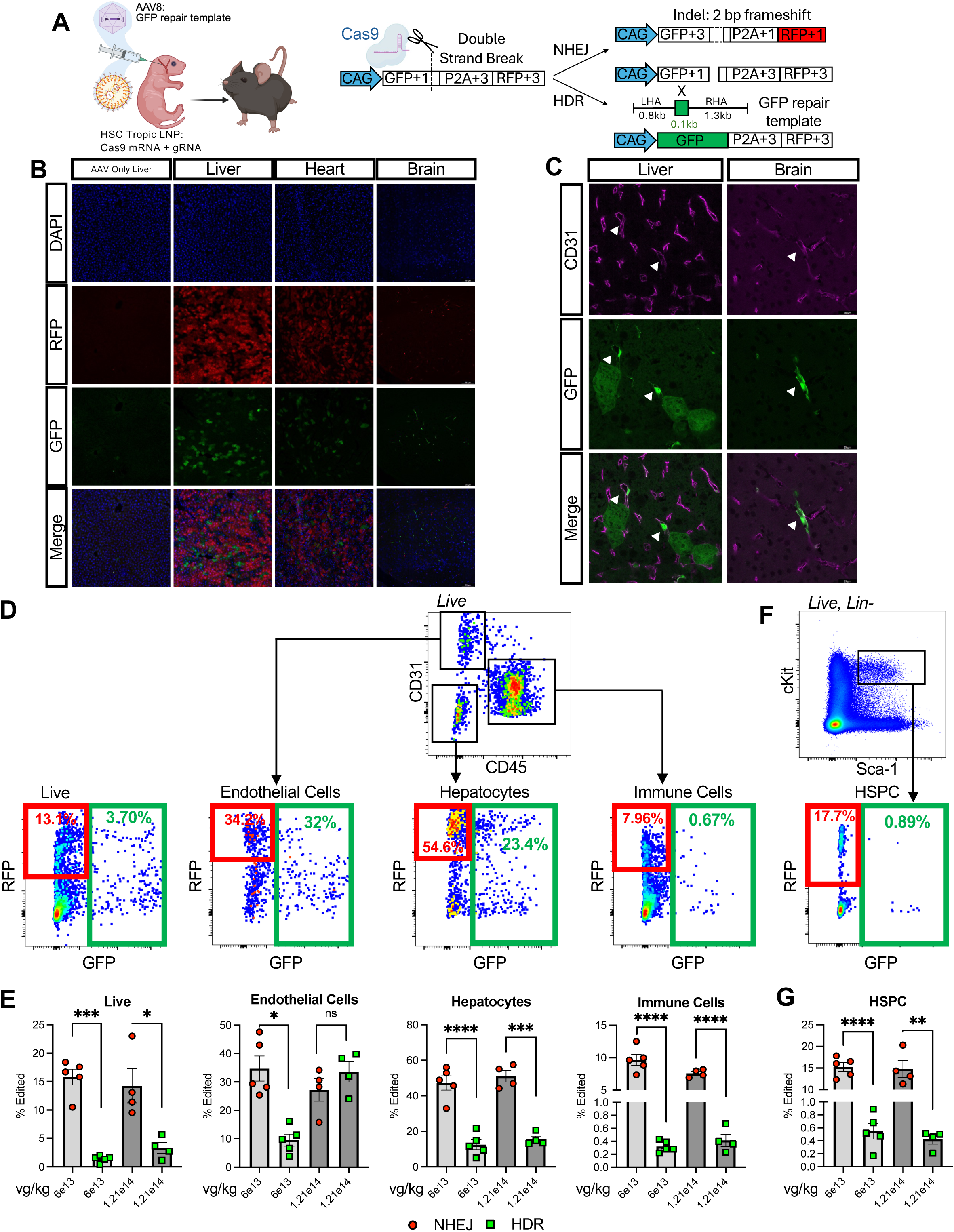
Combined LNP and AAV delivery enables *in vivo* homology-directed repair. **A)** Experiment schematic. P0 TLR mice were injected via the temporal vein with HSC-tropic LNP#1-Cas9/gRNA and AAV8-GFP repair. Liver and bone marrow were harvested in adult mice for editing analysis. **B)** Representative images of GFP expression (green) indicating HDR editing and RFP expression (red) indicating NHEJ editing in liver, heart and brain cells. **C)** Representative images of GFP expression (green) and CD31 (pink) in liver and brain, (original magnification x40). Arrows identify GFP+ cells co-stained with CD31. **D)** Gating strategy to identify GFP+ and RFP+ endothelial cells (CD31+, CD45-), hepatocytes (CD31-, CD45-) and immune cells (CD31-, CD45+) in the liver. **E)** Percentage of NHEJ (as measured by RFP+, red circles) and HDR (as measured by GFP+, green squares) edited cells at either 6e13 vg/kg or 1.21e14 vg/kg AAV8-GFP repair template. Differences in NHEJ vs HDR % editing at each dose measured by paired Student T-test. **F)** Gating strategy to identify GFP+ and RFP+ HSPCs (Lin-cKit+Sca+) in the bone marrow. **G)** Percentage of NHEJ (as measured by RFP+, red circles) and HDR (as measured by GFP+, green squares) edited HSPC at either 6e13 vg/kg or 1.21e14 vg/kg AAV8-GFP repair template. Differences in NHEJ vs HDR % editing at each dose measured by paired Student T-test. Data are mean+/-SEM; each dot represents an individual animal.*P<0.05, **P<0.01, ***P<0.001, P<0.0001

We designed a GFP restoration donor template to be packaged into an adeno-associated virus (AAV) vector and co-delivered with HSC-tropic LNP–Cas9/gRNA. Prior to *in vivo* experiments, we validated the repair template *in vitro* by electroporating Cas9/gRNA ribonucleoproteins into c-Kit–enriched bone marrow cells from TLR mice and transducing cells with AAV6 carrying the repair template. These experiments confirmed that the donor template supports GFP restoration and that AAV delivery alone does not produce GFP expression (**Fig. S4**).

We next injected neonatal (P0) TLR mice with HSC-tropic LNP-Cas9/gRNA together with AAV8 carrying the GFP repair template at varying vector genome doses. Tissues were analyzed 6-8 weeks after injection. Fluorescence imaging revealed GFP-positive cells in several tissues, including liver, heart and brain, indicating successful HDR-mediated repair *in vivo* (**Fig. 6B**). In the brain, GFP-positive cells were primarily CD31⁺ endothelial cells (**Fig. 6C**).

To quantify editing outcomes across liver cell populations, we performed flow cytometry to measure both NHEJ (RFP⁺) and HDR (GFP⁺) events in hepatocytes, endothelial cells and immune cells. As expected, NHEJ occurred at higher frequencies than HDR in most cell types (**Fig. 6D, E** and **Fig. S4B**). However, endothelial cells in the liver exhibited relatively robust HDR at higher AAV doses. Increasing AAV vector dose did not substantially increase HDR in other liver cell populations.

Finally, we examined editing outcomes in the bone marrow. Flow cytometric analysis confirmed robust NHEJ editing in hematopoietic stem and progenitor cells, consistent with earlier experiments. Notably, we also detected small but distinct populations of GFP⁺ HSPCs, indicating that HDR can occur in hematopoietic cells *in vivo* despite the limited tropism of AAV8 for these cell types (**Fig. 6F, G** and **Fig. S4C**). Together, these findings demonstrate that LNP-mediated delivery of Cas9 combined with viral delivery of a repair template can enable *in vivo* homology-directed repair across multiple tissues, including hematopoietic cells.

## Conclusion

In this study, we demonstrate that developmental hematopoiesis can be leveraged to enable efficient *in vivo* genome editing of HSCs. By exploiting the transient residence of HSCs in the fetal liver, we show that liver-tropic LNPs can access and edit bona fide long-term repopulating HSCs following in utero delivery. Edited cells retain their capacity for multilineage hematopoietic reconstitution and serial transplantation, confirming that genome editing does not impair stem cell function. In addition, we identify an LNP formulation that enhance HSC targeting during neonatal delivery, establishing that the window for *in vivo* HSC editing extends beyond fetal development into early postnatal life.

Our results further demonstrate that the early neonatal period, in which HSCs are circulating, remains permissive for *in vivo* HSC editing, particularly when using an optimized LNP formulation. Human newborns similarly exhibit circulating HSCs, suggesting that strategies combining prenatal diagnosis with early postnatal treatment could allow genome editing without bone marrow mobilization or antibody-mediated targeting approaches. Previous studies have demonstrated that mobilization of adult HSCs can improve editing efficiency^35,36^, highlighting the importance of stem cell accessibility for delivery platforms. HSCs are perhaps most accessible during fetal development, when they reside within the liver and LNP formulations that normally accumulate in the liver can access HSCs without the need for antibody-mediated targeting or stem cell mobilization.

Our strategy may have particular relevance for the treatment of severe congenital hematologic diseases. Many such disorders are now diagnosed prenatally through carrier screening, chorionic villus sampling, amniocentesis or noninvasive prenatal testing^23^. Consistently, the increase in clinical experience with intrauterine transfusions to treat diseases like alpha- thalassemia major^24^ has demonstrated the feasibility of delivering therapeutics during early development. Genome editing during fetal development or early neonatal life could therefore offer the possibility of correcting disease-causing mutations before irreversible pathology develops.

Importantly, our results also demonstrate that clinically meaningful levels of HSC editing can be achieved without the need for targeting ligands. Even partial correction of HSC populations may be therapeutically sufficient for certain diseases. For example, in hemoglobinopathies, relatively modest levels of corrected HSCs can generate substantial numbers of healthy erythroid cells due to the survival advantage of corrected red blood cells^1–3^. An additional strategy to improve correction is to combine the edit with editing of the EpoR, to enable more RBC production from edited HSCs^7^. In addition, diseases in which corrected hematopoietic stem cells have a selective advantage, such as Fanconi anemia^37^, may be particularly well suited for early-life genome editing strategies. Indeed, we have demonstrated that wild-type cells that engraft after *in utero* transplantation increase over time; *in vivo*-corrected cells could similarly increase over time.

Beyond generating insertions or deletions through NHEJ, we further demonstrate the feasibility of HDR *in vivo* using combined LNP delivery of Cas9 and viral delivery of a repair template. Although HDR efficiencies remain lower than those achieved through NHEJ, these findings highlight the potential to correct disease-causing mutations directly rather than relying solely on disruption strategies. Such approaches could circumvent the regulatory and manufacturing challenges for diseases in which patients carry diverse variants that otherwise require bespoke approaches. The levels of HDR achieved in the liver in this study (approximately 10% in hepatocytes and up to 30% in endothelial cells) may be clinically meaningful in multiple disease contexts. In particular, lysosomal storage diseases represent a promising target, as many are diagnosed prenatally and are characterized by early-onset pathology that may be mitigated by early intervention, including emerging *in utero* enzyme replacement strategies^29^. Moreover, editing of central nervous system endothelial cells could enable delivery of secreted proteins across the blood-brain barrier via cross-correction mechanisms, potentially extending the impact of systemic delivery. Notably, the efficiency of HDR in this model is likely limited by the tropism of the AAV vector used to deliver the repair template. Accordingly, the HDR efficiencies reported here likely underestimate the efficiency that could be achieved using donor delivery systems optimized for hematopoietic cells.

Safety considerations are particularly important in the context of fetal therapy, where both the fetus and the pregnant mother may be exposed to therapeutic agents. In our study, we did not observe evidence of overt maternal toxicity following *in utero* LNP exposure as assessed by maternal liver function tests. We also did not detect editing at predicted off-target sites using targeted sequencing approaches. Nevertheless, comprehensive safety evaluations such as including assessment of clonal hematopoiesis, rare genotoxic events, pharmacokinetic evaluations and lipid biodistribution studies in appropriate large animal models will be essential prior to clinical translation.

Together, these results provide proof-of-concept that the fetal and early neonatal periods represent a unique therapeutic window for *in vivo* genome editing of hematopoietic stem cells. By leveraging developmental hematopoiesis and scalable LNP delivery systems, this approach may enable one-time genome editing therapies for severe early-onset hematologic diseases. As prenatal diagnosis and fetal interventions continue to expand, early-life genome editing may constitute an important future strategy for treating genetic diseases at their earliest stages.

## Methods

### Mouse Husbandry and Injections

Mouse husbandry and procedures were performed according to the UCSF Institutional Animal Care and Use Committee-approved protocol AN202125-00B. mTmG (strain #007676) and TLR (strain #034033) mice were purchased from Jackson (Jax) Laboratories. Prenatal intrahepatic injections were performed as previously described^38^. Neonatal temporal vein injections were performed as previously described^39^. *In utero* and postnatal injection were dosed with same mg/kg of LNP based on previously published embryonic weights of C57Bl6 mice^40^ and weights at birth.

### LNP^67^ Formulation

Cre recombinase mRNA was diluted in 25 mM sodium acetate buffer, while the remaining lipid components were diluted in 100% ethanol. The phases were then microfluidically mixed^41,42^ using NanoAssemblr Ignite (Precision Nanosystems) at a relative flow rate of 3:1 (RNA and lipid phases, respectively) and 12ml/min total flow rate. LNPs were diluted 1:42 in sterile 10mM Tris buffer and dialyzed by centrifugation in 100 kDa ultra centrifuge tubes (Amicon) at 4°C. Sucrose (NF, EP, JP, ChP, high purity, low endotoxin, Fisher Scientific) was dissolved in a sterile 10 mM Tris buffer at 60% (w/v). LNPs in 10 mM Tris buffer were mixed with 60% (w/v) sucrose solution at 5:1 (v/v LNP:sucrose) for the final concentration of sucrose at 10% (w/v). LNPs were then flash-frozen in liquid nitrogen. Frozen LNPs were stored at ≤ −80°C. To thaw, LNPs were incubated at room temperature for 20 min. The LNPs were sterile-filtered with 0.22-µm filter before administration.

### LNP-Cas9/gRNA Formulations

Both LNP-Cas9/gRNA contain four lipid components: an ionizable lipid (IL), DSPC (1,2-distearoyl-sn-glycero-3-phosphocholine), cholesterol and a PEG-lipid (DMG-PEG2000; 1,2-Dimyristoyl-sn-glycerol-polyethylene glycol). The ionizable lipid used in the liver tropic LNP was LP01^43^ with a molar ratio of 50/9/38/3 (IL/DSPC/Chol/PEG-lipid). The ionizable lipid used in HSC-tropic LNP is described elsewhere^44^ where the molar ratio is 50/10/38.5/1.5 (IL/DSPC/Chol/DMG-PEG2000).

Both LNP-Cas9/gRNA were formulated by first dissolving the four lipid components in 100% ethanol at the appropriate concentration to enable a final N/P (amine-to-phosphate) ratio of 6. RNA cargo, at a 1:2 weight ratio of sgRNA-to-mRNA, was prepared in an acidic buffer containing 25 mM citrate buffer (pH 5) and 100 mM NaCl at a total RNA concentration of 0.45 mg/mL. LNPs were formulated as previously described^45^ using controlled mixing. All LNPs were characterized using a Ribogreen assay to determine cargo concentration and encapsulation efficiencies^46^. Dynamic light scattering using a Malvern NanoSizer was used to measure LNP size and polydispersity (PDI). Both LNP-Cas9/gRNA had encapsulation efficiencies above 95% and Z-average diameters between 70-95 nm.

### Tissue harvesting

Mice were euthanized by CO_2_ inhalation and transcardially perfused with cold PBS. Livers, leg and hip bones were processed as previously described^47,48^. Briefly, bone marrow was isolated by crushing and filtering. Livers were minced into 2mm x 2mm pieces and digested in 1mg/mL collagenase IV for 45 minutes, then passaged through a 19 g needle ten times, and filtered through a 70 µm filter.

### Tissue Processing for Microscopy

Freshly harvested tissues were fixed in 4% paraformaldehyde for 24 hours and cryopreserved in 30% sucrose prior to embedding in OCT. Tissue sections (10–20 μm) were prepared using a cryostat and mounted on glass slides. Sections from LNP^67^ treated mice were imaged using a an ECHO microscope without z-stacks with appropriate excitation and emission filters for GFP, tdTomato and DAPI. Nuclei were counterstained with DAPI prior to imaging. Sections from LNP- Cas9/gRNA treated mice were imaged using a Leica Stellaris confocal microscope without z-stacks (except for brains that were imaged with z-stack) with appropriate excitation and emission filters for GFP, RFP and DAPI.

For CD31 staining, tissue sections were permeabilized with 0.1% Triton X-100 and blocked with 5% normal serum. Sections were incubated with primary antibodies against CD31 (Fisher Scientific, AF3628, 1:200 dilution) overnight at 4 °C, followed by 1 hour incubation with donkey anti goat 633 (Biotium, 120127-1, 1:200 dilution). Nuclei were counterstained with DAPI prior to imaging. Imaging was performed on Leica Stellaris confocal microscope without z stacks. Images were processed using ImageJ or Fiji software.

### Maternal Liver Function Tests

Five to six days after *in utero* LNP^67^ injection, 500uL of peripheral blood was collected from injected dams via cardiac puncture following euthansia. Maternal blood samples were centrifuged at 14000 rpm for 10 minutes. The clear serum was aliquoted to a labeled Eppendorf tube. Serum samples were stored at −80 and sent for laboratory testing to measure serum liver function values at Zuckerberg San Francisco General Hospital Clinical Lab.

### Flow cytometry and FACS

Cell labeling was performed on ice in 1× PBS with 5 mM EDTA and 2% serum. Antibodies used are listed in Table S1 below. Analysis and cell sorting was performed on BD LSRII and BD FACS Aria II and analyzed using FlowJo. Cells were defined as follows as previously reported^47,49^: Granulocytes/Macrophages (Gr-1+ Mac-1+ B220- CD3- Ter119- CD61-); B cells (B220+ CD3- Gr-1- Mac-1- Ter119- CD61-); T cells (CD3+ B220- Gr-1- Mac-1- Ter119- CD61-); fetal/neonatal HSCs (Lin-cKit+Sca+SLAM+), adult HSC (Lin-cKit+Sca+SLAMmid/hi Flk2-), adult MPP (Lin-cKit+Sca+SLAM- Flk2+), HSPC (Lin-cKit+Sca+).

| Antibody | Source | Identifier | Dilution |
| --- | --- | --- | --- |
| B220-APC Cy7 | Biolegend | 103224 | 1:400 |
| CD3-A700 | Biolegend | 100216 | 1:200 |
| Mac-1-PE Cy7 | Biolegend | 101216 | 1:400 |
| Gr-1-Pacific Blue | Biolegend | 108430 | 1:400 |
| Ter119-PE Cy5 | Biolegend | 116210 | 1:400 |
| CD61-A647 | Biolegend | 104314 | 1:400 |
| B220-A700 | Biolegend | 103231 | 1:400 |
| Mac-1-A700 | Biolegend | 101222 | 1:400 |
| Gr-1-A700 | Biolegend | 108421 | 1:400 |
| Ter119-A700 | Biolegend | 116220 | 1:400 |
| cKit-APC Cy7 | Biolegend | 105826 | 1:400 |
| Sca-1-Pacific Blue | Biolegend | 122520 | 1:400 |
| SLAM-PE Cy7 | Biolegend | 115914 | 1:400 |
| Flk2-APC | Biolegend | 135310 | 1:100 |

### Transplantation experiments

Transplantation assays were performed as previously described^47^. Briefly, single purity sorted HSCs or HSPCs were isolated from liver or bone marrow of LNP-injected mice. Lineage markers were CD3, B220, Mac1, Gr1 and Ter119. Mac1 antibodies were excluded from the lineage cocktail when sorting fetal HSCs. Recipient mice aged 8-12 weeks were sublethally irradiated (500 rad for HSC transplants and 700 rad for HSPC transplants, single dose) with a Faxitron CP-160 and sorted cells (200 Tom+ or GFP+ HSCs or 1,000 RFP+ HSPCs) were transplanted retro-orbitally. Donor chimerism in peripheral blood of recipients was analyzed by flow cytometry. For secondary transplants, recipient mice aged 8-12 weeks were lethally irradiated (900 rad split dose) and transplanted with 10e6 whole bone marrow cells from primary recipients, as previously described^49^.

### Polymerase Chain Reaction (PCR) and Sanger sequencing

Genomic DNA was extracted from snap-frozen tissues using DNeasy Blood and Tissue kit (QIAGEN, catalog #69504) per manufacturer’s instructions, quantified using a nanodrop, and amplified using Phusion Green PCR MasterMix (ThermoFisher, catalog #F-566L) and the primers outline in Table S2. Thermocycler (Bio-Rad) settings were as follows: 98 °C (30 s), 98 °C (10 s), 72° C (30 s), 72 °C (30 s), return to step 2 for 34 cycles and then 72 °C (10 s). PCR products were purified using SPRIselect Beads (Beckman-Coulter, catalog #B23317) per manufacturer’s instructions, re-quantified using a nanodrop, and sent for sequencing (QuintaraBio). Sequencing (.ab1) files were analyzed using Synthego ICE software to determine indel percentage.

### Primer design for off-target amplicon sequencing

Off-target (OT) sites were first identified based on sequence homology to the on-target guide RNA and the presence of a compatible PAM. For each confirmed OT locus, primer pairs were designed to flank and fully encompass the predicted editing site, ensuring that the editing window was centrally located within the amplicon. Primers were designed to generate short PCR products of approximately 250–300 bp, which are optimal for high-quality amplicon-based next-generation sequencing and maximize coverage uniformity and sequencing accuracy. All primers were checked to avoid secondary structures, primer–dimer formation, and repetitive genomic regions, and to ensure unique genomic alignment. PCR amplification of each OT site was performed using these locus-specific primers, and the resulting amplicons were subjected to targeted deep sequencing using the Amplicon-EZ service (Azenta). This design strategy allowed sensitive and quantitative detection of rare editing events at candidate off-target loci.

### Off-target amplicon sequencing analysis

Raw amplicon-sequencing reads were first demultiplexed based on sample-specific barcodes to generate individual FASTQ files for each condition. The quality of the demultiplexed reads was assessed using FastQC, and the resulting reports were aggregated with MultiQC to evaluate overall sequencing quality, including base quality distributions, GC content, adapter contamination, and consistency across samples. Only datasets passing quality control were used for downstream analysis. Editing outcomes at candidate off-target sites were quantified using CRISPRessoBatch. A quantification window of 20 bp centered on the predicted Cas9 cleavage site was used to determine indel frequencies. Editing frequencies were compared with untreated controls to determine whether editing exceeded the assay’s limit of detection.

### TIDE analysis

Editing efficiencies from Sanger sequencing traces were quantified using the Tracking of InDels by DEcomposition (TIDE) algorithm. Sequencing chromatograms were analyzed using the TIDE web tool (version 3.3.0) with default parameters to estimate insertion and deletion frequencies as previously described^50^.

### AAV6 vector for *in vitro* HDR validation

Adeno-associated virus (AAV) vector plasmids were cloned into the pAAV-MCS plasmid, comprised of inverted terminal repeats (ITRs) derived from AAV2. Gibson Assembly Mastermix (New England Biolabs, Ipswich, MA) was used for the creation of all DNA repair vectors according to manufacturer’s instructions. Once cloned, plasmids were transformed into Stable Competent E. coli (New England Biolabs) and amplified using Plasmid Plus Midi Kits (QIAGEN N.V., Hilden, Germany). For *in vitro* testing, AAV6 vector was produced with little variation as previously described^51^. 293T cells (ATCC, Manassas, VA) were seeded in five 15cm^2^ dishes with 1.3–1.5×10^7^ cells per plate 24h pre-transfection. Each dish was then transfected with 112μg polyethylenimine, 6μg ITR-containing plasmid, and 22μg pDGM6 (gift from David Russell, University of Washington), which expresses AAV6 cap, AAV2 rep, and Ad5 helper genes. Following a 48-72h incubation, cells were harvested, and vectors were purified using the Takara Bio AAVpro purification kit as per manufacturer’s instructions and stored at −80°C until further use. AAV6 vectors were titered using a Bio-Rad QX200 ddPCR machine and QuantaSoft software (v.1.7) to measure the number of vector genomes, as described previously^52^.

### AAV8 vector packaging for *in vivo* HDR experiments

Recombinant adeno-associated virus (rAAV) was produced by SignaGen Laboratories (Frederick, Maryland) using a helper-free transient transfection system in HEK293-derived packaging cells. The production process involved co-transfection of three plasmids: (i) the AAV cis plasmid containing the transgene expression cassette flanked by AAV inverted terminal repeats (ITRs), (ii) a packaging plasmid encoding the AAV Rep and Cap proteins corresponding to the desired serotype, and (iii) a helper plasmid supplying essential adenoviral gene products required for efficient AAV replication and assembly. Following transfection, cells were maintained under optimized culture conditions to facilitate viral production. At approximately 72 hours post-transfection, cells and culture supernatant were harvested. Viral particles were released via cell lysis and subsequently purified by cesium chloride (CsCl) gradient ultracentrifugation to achieve high-purity vector preparations. The purified vector was then formulated in an appropriate buffer for research use. Vector genome (VG) titers were determined by quantitative polymerase chain reaction (qPCR) using primers specific to the AAV ITR region. Prior to analysis, samples were treated with DNase to eliminate residual plasmid DNA, and encapsidated viral genomes were subsequently released via proteinase digestion and heat denaturation. Quantification was performed against a standard curve generated from a linearized DNA template containing the ITR target sequence. Final titers were reported as vector genomes per milliliter (vg/mL).

### Statistical analysis

Results were graphed and analyzed using statistical software GraphPad Prism version 9.5.1 (GraphPad Software). Comparisons between two groups were performed using Student’s t-test or Mann–Whitney test as indicated. Multiple comparisons were analyzed using one-way ANOVA with Tukey’s correction. P < 0.05 was considered statistically significant.

## Acknowledgements

This work was supported by NIH R01 HL161291 to T.C.M. and UCSF CIRM Postdoctoral Training grant EDUC4-12812 to A.K.W. We acknowledge the UCSF Parnassus Flow Cytometry Core (RRID:*SCR_018206*) supported in part by Grant NIH P30 DK063720 and by the NIH S10 Instrumentation Grant S10 1S10OD021822-01. We thank all the members of the MacKenzie lab, Dahlman lab and Cromer lab for helpful discussion.

## Author contributions

A.K.W. and T.C.M conceptualized the study. A.K.W, B.B., T.L., F.M.Z., M.A.C., A.D., A.G., B.P., D.O., K.J. performed experiments. E.S.E., R.Z., O.C., H.K., R. M., C.S., P.G.M., M.O., C.B., S.B. and J.E.D. formulated the LNPs. T.M. and H.S. generated AAV for *in vitro* experiments. A.K.W., B.B., A.D. and M.K.C. analyzed the data. A.K.W. prepared the figures. A.K.W. and T.C.M wrote the manuscript with input from all authors.

## Figure Legends

**Figure S1.**
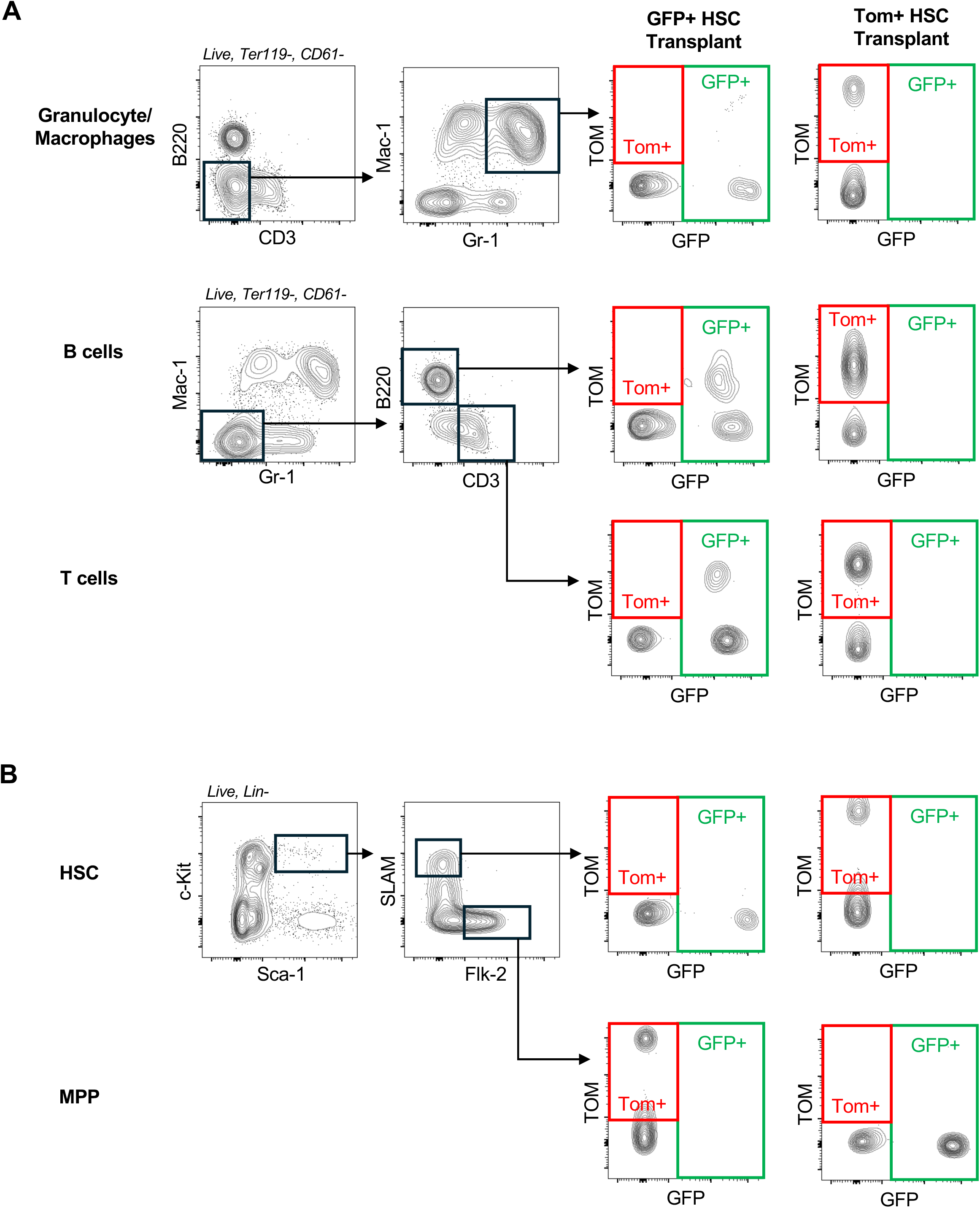
LNP^67^ mTmG HSC transplant gating strategy. **A)** Representative FACS plots of granulocyte/macrophage, B and T cell gating to identify GFP+ HSC derived and Tom+ HSC derived mature cells in the peripheral blood of transplanted mice. **B)** Representative FACS plots of HSCs and MPPs too identify GFP+ HSC derived and Tom+ HSC derived HSPCs in the bone marrow of transplanted mice.

**Figure S2.**
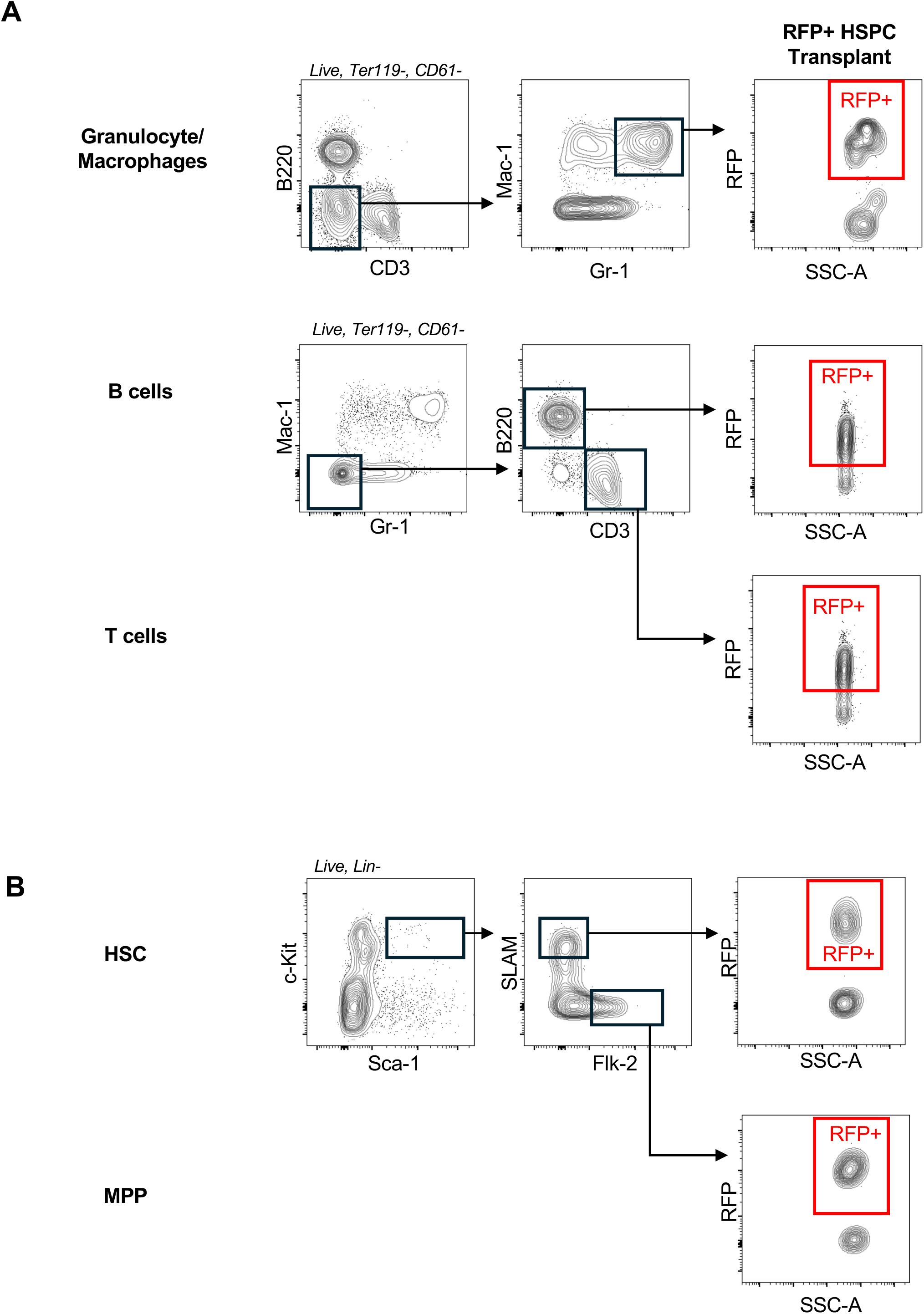
Liver-tropic LNP TLR HSCP transplant gating strategy. **A)** Representative FACS plots of granulocyte/macrophage, B and T cell gating to identify RFP+ HSC derived mature cells in the peripheral blood of transplanted mice. **B)** Representative FACS plots of HSCs and MPPs too identify RFP+ HSC derived HSPCs in the bone marrow of transplanted mice.

**Figure S3.**
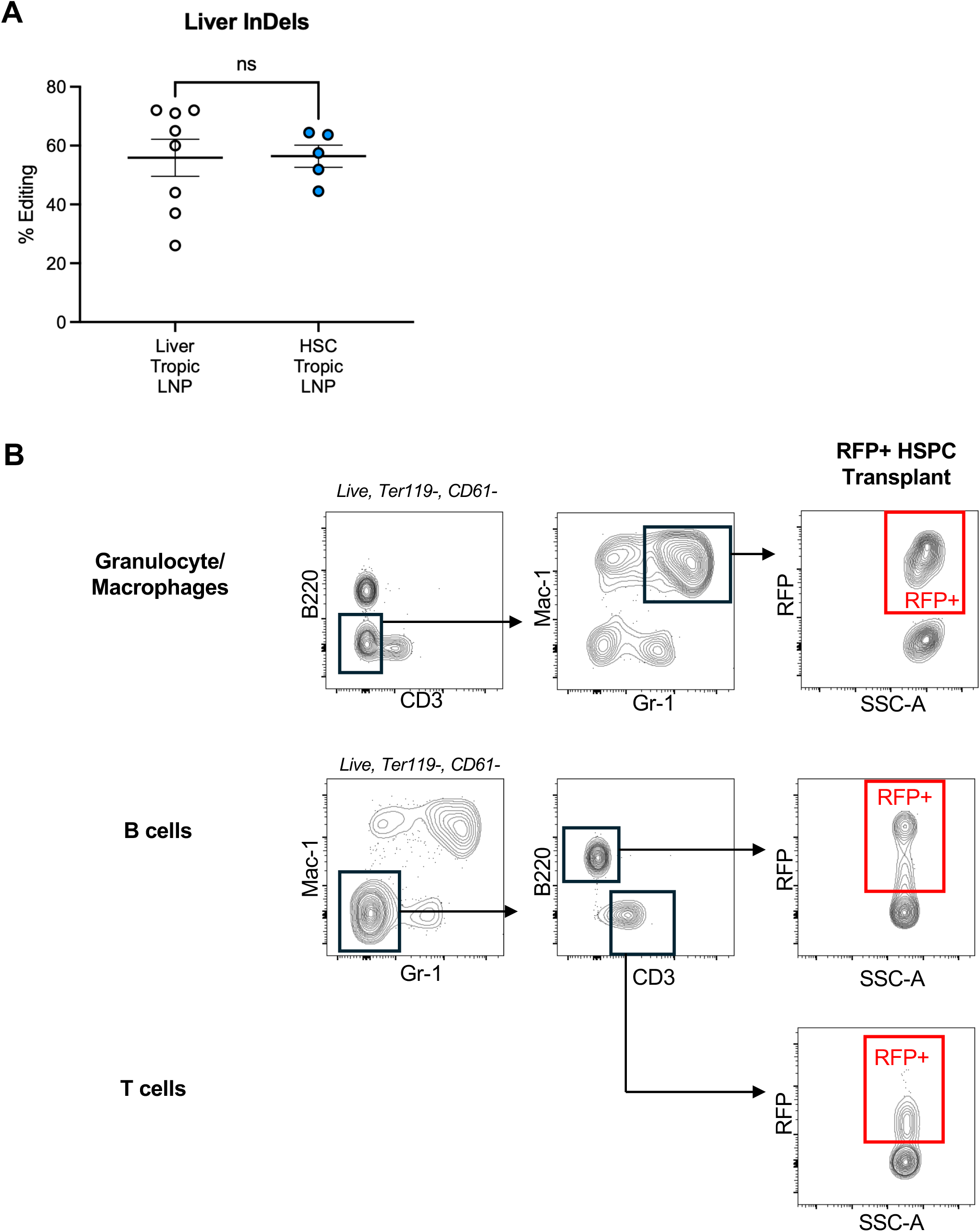
HSC-tropic LNP liver editing and TLR HSPC transplant gating strategy. **A)** Representative FACS plots of granulocyte/macrophage, B and T cell gating to identify RFP+ HSC derived mature cells in the peripheral blood of transplanted mice. **B)** Percentage of InDels by Sanger sequencing in the liver of mice injected postnatally with 3 mg/kg of different LNP formulations. P=NS (One-way ANOVA with Tukey’s multiple comparisons test).

**Figure S4.**
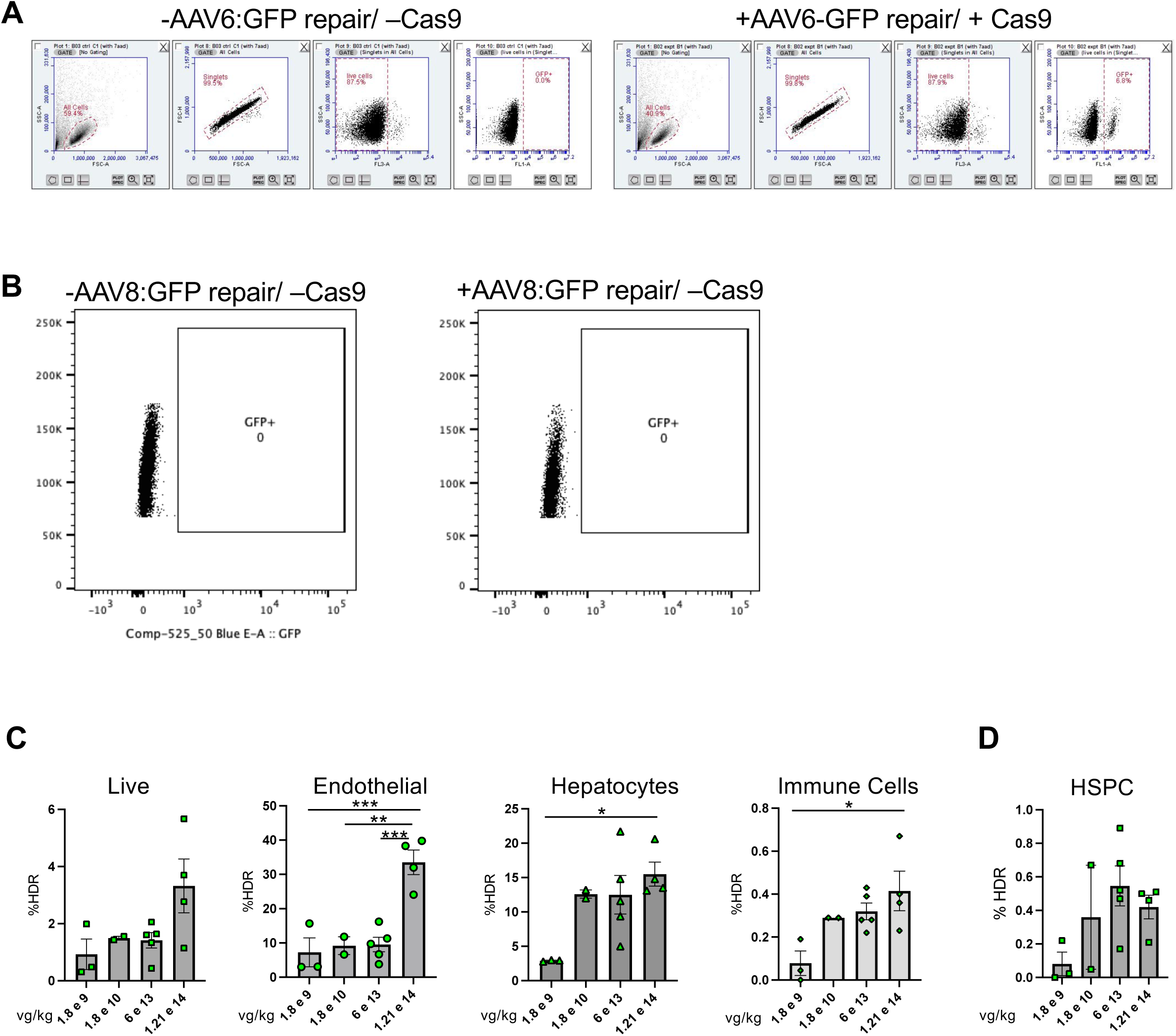
*In Vitro* AAV-GFP validation and AAV8 dose response. **A)** Representative FACS plots of GFP expression in TLR cKit enriched bone marrow cells in culture electroporated with the Cas9/gRNA RNP and transduced with and without AAV6+GFP repair template. **B)** Percentage of HDR (as measured by GFP) edited liver cell populations at 1.8e9, 1.8e10, 6e13 or 1.21e14 vg/kg AAV8-GFP repair template. Data are mean+/-SEM; each dot represents an individual animal. Differences measure by One-way ANOVA with Tukey’s multiple comparisons test, *P<0.05 **P<0.01, ***P<0.001)

**Table S1.**
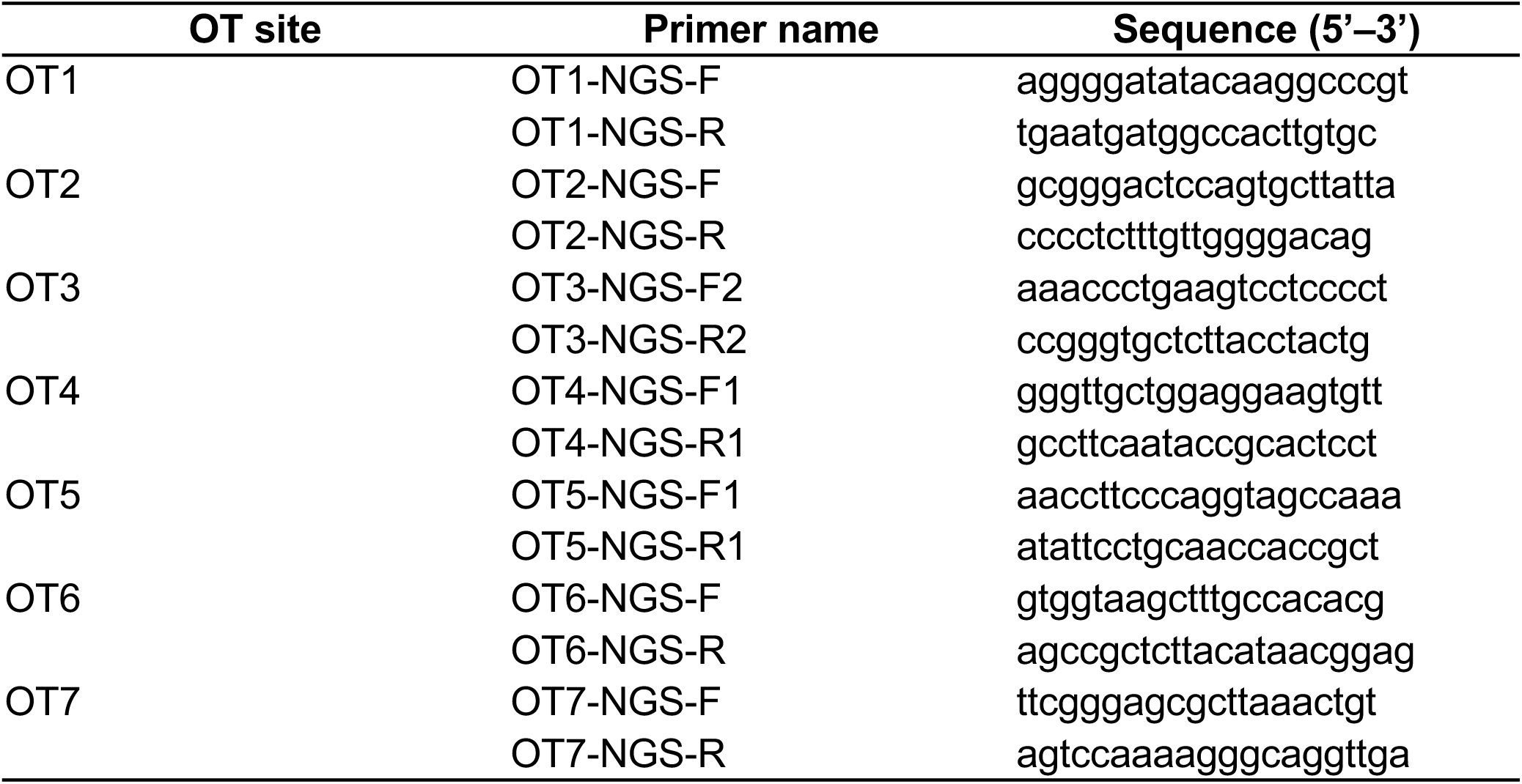
Primer sequences used for off-target (OT) amplicon sequencing.

**Table S2.**
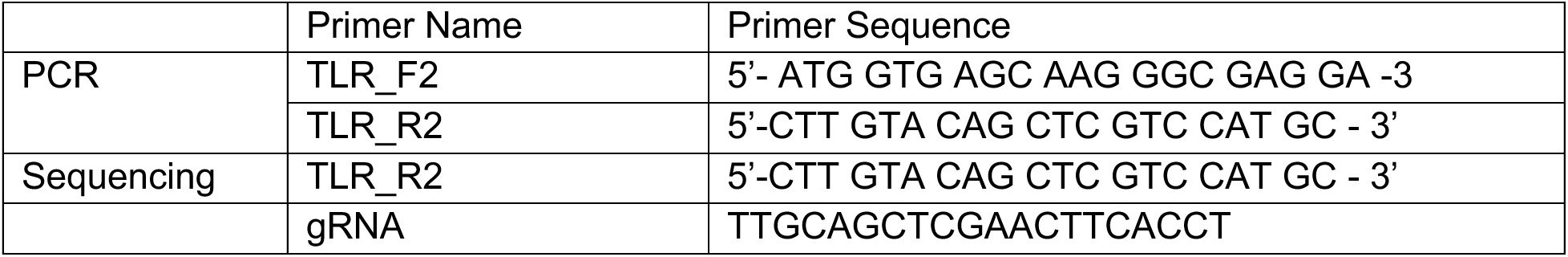
Primers used for PCR and Sanger Sequencing.

## References

1. Fitzhugh, C. D. et al. At least 20% donor myeloid chimerism is necessary to reverse the sickle phenotype after allogeneic HSCT. Blood 130, 1946–1948 (2017).

2. Chaudhury, S. et al. A Multicenter Retrospective Analysis Stressing the Importance of Long-Term Follow-Up after Hematopoietic Cell Transplantation for β-Thalassemia. Biol Blood Marrow Transplant 23, 1695–1700 (2017).

3. Hsieh, M. M., Wu, C. J. & Tisdale, J. F. In mixed hematopoietic chimerism, the donor red cells win. Haematologica 96, 13–15 (2011).

4. Frangoul, H. et al. CRISPR-Cas9 Gene Editing for Sickle Cell Disease and β- Thalassemia. The New England journal of medicine 384, 252–260 (2021).

5. Kanter, J. et al. Biologic and Clinical Efficacy of LentiGlobin for Sickle Cell Disease. N Engl J Med 386, 617–628 (2022).

6. Segura, E. E. R. et al. Lentiviral vectors for hematopoietic stem cell gene therapy restore α-globin expression in α-thalassemia red blood cells. Cell Rep Med 6, 102362 (2025).

7. Chu, S. N. et al. Dual α-globin-truncated erythropoietin receptor knockin restores hemoglobin production in α-thalassemia-derived erythroid cells. Cell Rep 44, 115141 (2025).

8. Charlesworth, C. T., Hsu, I., Wilkinson, A. C. & Nakauchi, H. Immunological barriers to haematopoietic stem cell gene therapy. Nat Rev Immunol 22, 719–733 (2022).

9. Rueda, J., de Miguel Beriain, Í. & Montoliu, L. Affordable Pricing of CRISPR Treatments is a Pressing Ethical Imperative. CRISPR J 7, 220–226 (2024).

10. Sheridan, C. The world’s first CRISPR therapy is approved: who will receive it? Nature Biotechnology 42, 3–4 (2023).

11. Hou, X., Zaks, T., Langer, R. & Dong, Y. Lipid nanoparticles for mRNA delivery. Nat Rev Mater 6, 1078–1094 (2021).

12. Polack, F. P. et al. Safety and Efficacy of the BNT162b2 mRNA Covid-19 Vaccine. N Engl J Med 383, 2603–2615 (2020).

13. Baden, L. R. et al. Efficacy and Safety of the mRNA-1273 SARS-CoV-2 Vaccine. N Engl J Med 384, 403–416 (2021).

14. Gillmore, J. D. et al. CRISPR-Cas9 In Vivo Gene Editing for Transthyretin Amyloidosis. N Engl J Med 385, 493–502 (2021).

15. Musunuru, K. et al. Patient-Specific In Vivo Gene Editing to Treat a Rare Genetic Disease. New England Journal of Medicine 392, 2235–2243 (2025).

16. Hosseini-Kharat, M., Bremmell, K. E. & Prestidge, C. A. Why do lipid nanoparticles target the liver? Understanding of biodistribution and liver-specific tropism. Mol Ther Methods Clin Dev 33, 101436 (2025).

17. Breda, L. et al. In vivo hematopoietic stem cell modification by mRNA delivery. Science 381, 436–443 (2023).

18. Palanki, R. et al. In utero delivery of targeted ionizable lipid nanoparticles facilitates in vivo gene editing of hematopoietic stem cells. Proc Natl Acad Sci U S A 121, e2400783121 (2024).

19. Cain, T. L., Derecka, M. & McKinney-Freeman, S. The role of the haematopoietic stem cell niche in development and ageing. Nat Rev Mol Cell Biol 26, 32–50 (2025).

20. Morrison, S. J., Hemmati, H. D., Wandycz, A. M. & Weissman, I. L. The purification and characterization of fetal liver hematopoietic stem cells. Proceedings of the National Academy of Sciences of the United States of America 92, 10302–6 (1995).

21. Bowie, M. B. et al. Hematopoietic stem cells proliferate until after birth and show a reversible phase-specific engraftment defect. J Clin Invest 116, 2808–2816 (2006).

22. Lin, S., Staahl, B. T., Alla, R. K. & Doudna, J. A. Enhanced homology-directed human genome engineering by controlled timing of CRISPR/Cas9 delivery. Elife 3, e04766 (2014).

23. Shear, M. A. et al. The Expanding Role of Gene Sequencing in Shaping Fetal Therapies: Clinical and Ethical Considerations. Prenat Diagn https://doi.org/10.1002/pd.6890 (2025) doi:10.1002/pd.6890.

24. Schwab, M. E. et al. The impact of in utero transfusions on perinatal outcomes in patients with alpha thalassemia major: the UCSF registry. Blood Adv 7, 269–279 (2023).

25. Derderian, S. C., Jeanty, C., Walters, M. C., Vichinsky, E. & MacKenzie, T. C. In utero hematopoietic cell transplantation for hemoglobinopathies. Front Pharmacol 5, 278 (2014).

26. Flake, A. W. et al. Treatment of X-linked severe combined immunodeficiency by in utero transplantation of paternal bone marrow. N Engl J Med 335, 1806–1810 (1996).

27. MacKenzie, T. C. et al. In Utero Hematopoietic Cell Transplantation in Fetuses with Alpha Thalassemia Major: A Phase 1 Clinical Trial. Blood Adv bloodadvances.2026019655 (2026) doi:10.1182/bloodadvances.2026019655.

28. Cohen, J. L. et al. In Utero Enzyme-Replacement Therapy for Infantile-Onset Pompe’s Disease. N Engl J Med 387, 2150–2158 (2022).

29. Herzeg, A. et al. Intrauterine enzyme replacement therapies for lysosomal storage disorders: Current developments and promising future prospects. Prenat Diagn 43, 1638–1649 (2023).

30. Schwab, M. E. & MacKenzie, T. C. Prenatal Gene Therapy. Clin Obstet Gynecol 64, 876–885 (2021).

31. Christensen, J. L., Wright, D. E., Wagers, A. J. & Weissman, I. L. Circulation and chemotaxis of fetal hematopoietic stem cells. PLoS Biol 2, E75 (2004).

32. Kim, H. et al. Lipid nanoparticle-mediated mRNA delivery to CD34+ cells in rhesus monkeys. Nat Biotechnol https://doi.org/10.1038/s41587-024-02470-2 (2024) doi:10.1038/s41587-024-02470-2.

33. Muzumdar, M. D., Tasic, B., Miyamichi, K., Li, L. & Luo, L. A global double-fluorescent Cre reporter mouse. Genesis (New York, N.Y. : 2000) 45, 593–605 (2007).

34. Chu, V. T. et al. Increasing the efficiency of homology-directed repair for CRISPR- Cas9-induced precise gene editing in mammalian cells. Nat Biotechnol 33, 543–548 (2015).

35. Li, C. et al. In vivo HSC prime editing rescues sickle cell disease in a mouse model. Blood 141, 2085–2099 (2023).

36. Milani, M. et al. In vivo haemopoietic stem cell gene therapy enabled by postnatal trafficking. Nature 643, 1097–1106 (2025).

37. Swartzrock, L. et al. In utero hematopoietic stem cell transplantation for Fanconi anemia. Blood Adv 8, 4554–4558 (2024).

38. Nijagal, A., Le, T., Wegorzewska, M. & Mackenzie, T. C. A mouse model of in utero transplantation. J Vis Exp 2303 (2011) doi:10.3791/2303.

39. Gombash Lampe, S. E., Kaspar, B. K. & Foust, K. D. Intravenous injections in neonatal mice. J Vis Exp e52037 (2014) doi:10.3791/52037.

40. Kulandavelu, S. et al. Embryonic and neonatal phenotyping of genetically engineered mice. ILAR J 47, 103–117 (2006).

41. Chen, D. et al. Rapid discovery of potent siRNA-containing lipid nanoparticles enabled by controlled microfluidic formulation. J Am Chem Soc 134, 6948–6951 (2012).

42. Belliveau, N. M. et al. Microfluidic Synthesis of Highly Potent Limit-size Lipid Nanoparticles for In Vivo Delivery of siRNA. Mol Ther Nucleic Acids 1, e37 (2012).

43. Finn, J. D., et al. A Single Administration of CRISPR/Cas9 Lipid Nanoparticles Achieves Robust and Persistent In Vivo Genome Editing. Cell Rep 22, 2227–2235 (2018).

44 . Ionizable amine lipids and lipid nanoparticles - Patent WO-2020219876-A1 - PubChem. https://pubchem.ncbi.nlm.nih.gov/patent/WO-2020219876-A1.

45. Modified amine lipids - Patent WO-2020118041-A1 - PubChem. https://pubchem.ncbi.nlm.nih.gov/patent/WO-2020118041-A1.

46. Leung, A. K. K. et al. Lipid Nanoparticles Containing siRNA Synthesized by Microfluidic Mixing Exhibit an Electron-Dense Nanostructured Core. J Phys Chem C Nanomater Interfaces 116, 18440–18450 (2012).

47. Worthington, A. K., et al. IL7Rα, but not Flk2, is required for hematopoietic stem cell reconstitution of tissue-resident lymphoid cells. Development (Cambridge, England) 149, (2022).

48. Leung, G. A. et al. The lymphoid-associated interleukin 7 receptor (IL7R) regulates tissue-resident macrophage development. Development (Cambridge) 146, (2019).

49. Beaudin, A. E. et al. A Transient Developmental Hematopoietic Stem Cell Gives Rise to Innate-like B and T Cells. Cell stem cell 19, 768–783 (2016).

50. Brinkman, E. K., Chen, T., Amendola, M. & van Steensel, B. Easy quantitative assessment of genome editing by sequence trace decomposition. Nucleic Acids Res 42, e168 (2014).

51. Khan, I. F., Hirata, R. K. & Russell, D. W. AAV-mediated gene targeting methods for human cells. Nat Protoc 6, 482–501 (2011).

52. Aurnhammer, C. et al. Universal real-time PCR for the detection and quantification of adeno-associated virus serotype 2-derived inverted terminal repeat sequences. Hum Gene Ther Methods 23, 18–28 (2012).

